# Substrate profiling of human Transglutaminase 1 using cDNA display and next-generation sequencing

**DOI:** 10.1101/2023.10.22.563423

**Authors:** T.I.K. Munaweera, Jasmina Damnjanović, Maurizio Camagna, Moeri Nezu, Beixi Jia, Kiyotaka Hitomi, Naoto Nemoto, Hideo Nakano

## Abstract

Human transglutaminase 1 regulates skin development and is linked with several disease conditions with yet fully unknown mechanisms. To uncover all of its roles in health and disease, the understanding of protein substrates and their reactivity with transglutaminase 1 is necessary. To gain insight into the substrate profile of human transglutaminase 1, this study uses an *in vitro* selection system based on the cDNA display technology to screen two displayed peptide libraries differing in the number and position of mutated sites relative to the reactive glutamine, in terms of their reactivity to transglutaminase 1. Analysis of the selected DNA pools of by next-generation sequencing and in-house bioinformatics methods revealed a detailed transglutaminase 1 substrate profile indicating preferred and non-preferred amino acid sequences. We have identified a peptide with sequence AEQHKLPSKWPF showing high reactivity to human transglutaminase 1 and low reactivity to transglutaminase 2 and transglutaminase 3. The position weight matrix consisting of per residue amino acid enrichment factors of all selected peptides was used to search human proteins by our in-house search algorithm and identify highly scoring sequence motifs in their primary structure. The search identified six already known transglutaminase 1 substrate proteins as highly scored hits as well as a list of candidate substrates that are under investigation.

## Introduction

Mammalian transglutaminases (TGs) catalyze the formation of Ca^2+^-dependent Nε (γ-glutamyl) lysine isopeptide bond cross-linking among protein-bound glutamine (Gln) and lysine (Lys) residues (Eckert et al., 2015, Hitomi, Ed. 2015). The transamidation reaction catalyzed by transglutaminases proceeds by attaching a primary amine to the glutamine residue, while deamination occurs when the glutamine acceptor molecule is replaced with water to form glutamic acid (Savoca et al., 2018). TG family is composed of eight catalytically active isozymes (TG1-TG7, and Factor XIII), and an inactive erythrocyte protein band 4.2 (EPB4.2) (Savoca et al., 2018). These enzymes are involved in multiple biological processes such as the formation of the skin epidermis, and blood clotting, and their abnormal activities are found to be linked with various disease conditions (Eckert et al., 2005; Lorand & Graham, 2003; Mehta Kapil, 2005).

TG1 is expressed in the epithelial cells, namely in the spinous and granular layers of the epidermis. The barrier function in the outermost layer of the human skin epidermis is maintained by the cornified envelope (CE), formed beneath the plasma membrane in the terminally differentiated keratinocytes (Candi et al., 2005). The TGs 1, 2, 3, and 5 in the keratinocytes are responsible for cross-linking structural proteins such as loricrin, involucrin, and small proline-rich family with distinct preference to form CE (Hitomi, 2005). The membrane-bound TG1 is considered the most abundant isozyme that is predominantly involved in epithelial differentiation (Esposito & Caputo, 2005). The importance of TG1 is further revealed by the identification of human TG1 mutations related to congenital ichthyosis, including lamellar ichthyosis and non-bullous congenital ichthyosiform erythroderma (Huber et al., 1995; Hitomi, 2005; Sugimura et al., 2008a). Furthermore, differential expression of TG1 is suspected to be linked with Alzheimer’s disease (Tripathy et al., 2020).

Even though the reaction mechanism of TG isozymes follows the same pattern, they are known to have unique substrates that are involved in different biological events (Facchiano & Facchiano, 2005; Hitomi et al. Ed. 2015). Each isozyme is known to recognize different cross-linking sites in the substrate proteins implying that the primary/tertiary sequence surrounding the reactive Gln residue is the governing factor of the interaction between each isoenzyme and its substrate (Hitomi et al., 2009). Isozyme-specific substrates are used for designing artificial fluorescently labeled Gln probes that are being utilized for the detection of *in vitro* and *in situ* activity of TGs (Sugimura et al., 2006, 2008; Hitomi et al., 2009; Tatsukawa et al., 2022). Several isozyme-specific substrates of TG1 have been discovered previously by phage display screening, and among them, the K5 peptide with the amino acid sequence YEQHKLPSSWPF (Sugimura et al., 2008) has been found to exhibit the highest specificity and strong reactivity. Alongside K5, quite a few other peptides with consensus sequence QxR(K)LxxxWP showed high specificities and reactivities, indicating the importance of the conserved residues for reactivity with TG1. However, although this study provided important seeds and insights on the TG1 substrate preference, during the time it was done, technical limitations did not allow for a comprehensive analysis of the whole selected pool of peptide sequences, because the number of clones to analyze was limited by the available sequencing technologies and yet unavailable computational tools for big data analysis.

Aiming to comprehensively analyze the substrate preference of TG2, we have previously established a cDNA display-based selection system coupled with next-generation sequencing and bioinformatics for substrate profiling of this enzyme (Damnjanović et al., 2022). In the present study, the existing platform with changes in the library design and preparation, as well in the number of selection rounds, was used to study the substrate preference of TG1 by selection from the random libraries designed based on the existing K5 peptide backbone. The consensus sequence determined for K5 and other peptides selected using the phage display study was considered important for substrate reactivity with TG1 and was, thus, kept constant. The randomized DNA libraries were *in vitro* transcribed, displayed, translated, and used for TG1 activity-based selection. The selected libraries were analyzed by NGS and bioinformatics, which revealed several preferred as well as non-preferred sequences surrounding the reactive Gln residue, and a number of candidate Gln substrates with a higher enrichment factor than the known K5 sequence. The new candidate substrates were analyzed by *in vitro* TG enzymatic assays to verify their reactivity with isozyme specificity.

## MATERIALS AND METHODS

### Materials

Human keratinocyte transglutaminase (TG1), tissue transglutaminase (TG2), and epidermal transglutaminase (TG3) were obtained from Zedira GmbH (Darmstadt, Germany) as recombinant proteins produced and purified from infected insect cells or *E. coli*. Puromycin cnvK linker was obtained from Epsilon Molecular Engineering Inc. (Saitama, Japan). N-terminal biotinylated peptides were custom-synthesized by Pepmic (Suzhou, China). Oligonucleotide primers were ordered from Eurofins genomics (Tokyo, Japan), and are listed in Table S1.

### Preparation of DNA libraries

Two DNA libraries (Figure S1), Single Mutant Library (K5-SML) with a single mutation introduced by NNK codon in either of 8 positions of the K5 backbone, and the Semi Random Library (K5-SRL) with multiple mutations in 7 positions of the K5 backbone, were used for the selection of preferred substrate sequences in TG1-catalyzed enzymatic reaction. The mutated peptide genes were custom-ordered from Eurofins genomics (Tokyo, Japan) as single-stranded DNA of 78 bases containing the peptide coding region, as well as positions of DNA upstream and downstream of the peptide sequence necessary for the DNA library preparation. ssDNA fragments were transformed to dsDNA by Klenow fragment DNA polymerase (Takara Bio, Shiga, Japan) with primer 1, and assembled with PCR amplified (primers 2 and 3) pRSET vector fragment using Gibson assembly (NEB Japan, Tokyo, Japan) to add T7 promoter, T7 terminator, and sequences necessary for display formation to the peptide genes. This was followed by PCR amplification using primers 4 and 5 and column purification to prepare the initial DNA library ready for *in vitro* transcription.

### Preparation of cDNA display

cDNA displayed peptide libraries were prepared in the same manner as described previously (Damnajnovic et al., 2022) and shown in Figure 1A. In short, DNA libraries were used as templates for *in vitro* transcription by RiboMAX Large-scale RNA production system-T7 (Promega). The reaction mixture was incubated at 37°C for 2 h, followed by RNA purification using NucleoSpin RNA kit (Takara Bio, Shiga, Japan) with on-column DNA digestion. The quality of produced RNA was checked by absorbance using NanoDrop (Thermo Fisher) and Urea PAGE after staining with SYBR Gold (Invitrogen). Gels were visualized by an LED light imager (Biotools, Gunma, Japan) equipped with a green band-pass filter and imaging software MISVS II. Photo-crosslinking of mRNA and puromycin linker including cnvK (Mochizuki et al., 2015) was done on a 20-μL scale by hybridization of the mRNA to the linker followed by UV irradiation at 365 nm for 4 min. Confirmation of the crosslinking reaction was done by 8M Urea PAGE with two detection methods, fluorescent signal detection coming from the fluorescein attached to the linker, and staining with SYBR Gold. The gels were visualized with an LED light imager equipped with a green band-pass filter and imaging software MISVS II. The *in vitro* translation of peptides to synthesize mRNA display was performed with PURE*frex*2.1 (GeneFrontier, Kashiwa, Japan) on a 25-μL scale in triplicates. Incubation of the mixture at 37°C for 30 min was followed by 5 min incubation at 37°C with 20 mM EDTA to release the ribosomes. Confirmation of mRNA display formation was done by Urea SDS-PAGE with fluorescein detection. The visualization of gels was done using the same imaging system as for Urea PAGE. Triplicate reactions of mRNA display synthesis were joined, immobilized to the streptavidin-coated magnetic beads (Dynabeads MyOne Streptavidin C1, Invitrogen) and converted to cDNA display using ReverTraAce (Toyobo, Tokyo, Japan) reverse transcriptase at 42°C for 30 min. Treatment of the beads with RNaseT1 (Thermo Fisher) for 15 min at 37°C on a rotator was applied to release the formed cDNA display from the beads using the RNaseT1 recognition site enclosed in the puromycin linker. Collected supernatant containing mRNA-cDNA-linker-peptide complexes was used in the subsequent purification process.

**Figure 1.**
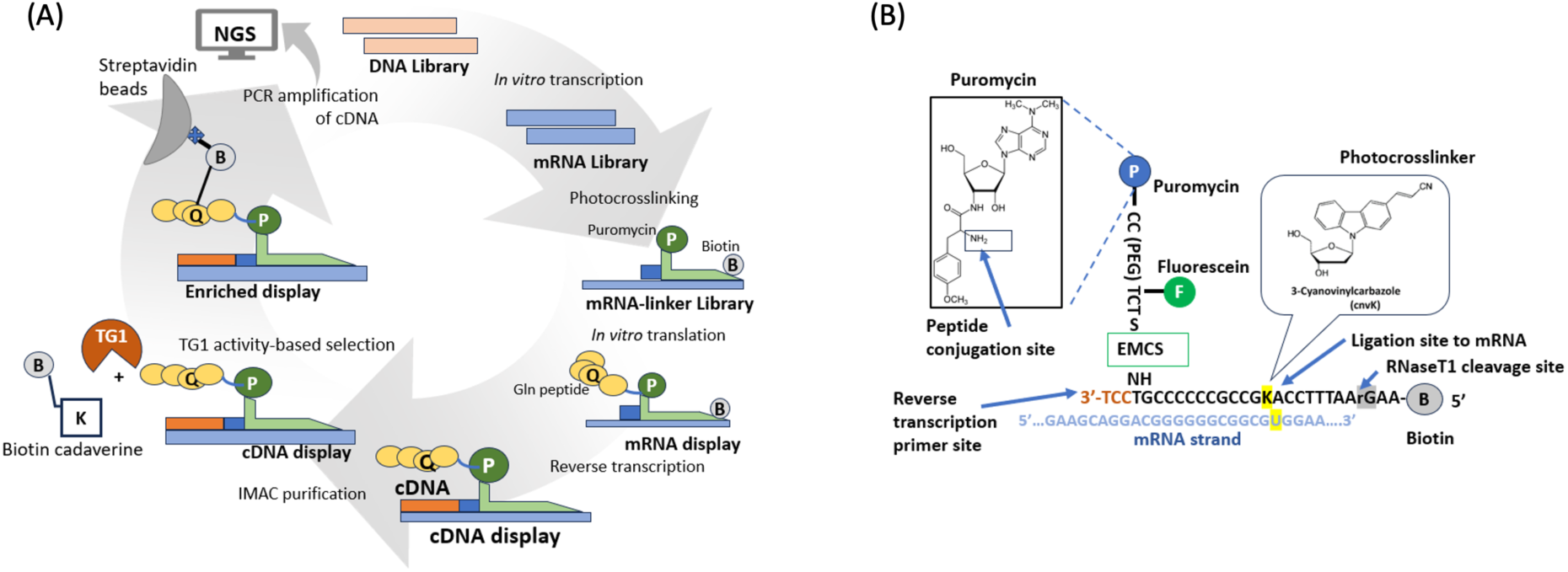
(A) Schematic diagram of the cDNA display platform used in this study; (B) Structure of the puromycin linker (adopted with modifications from Nemoto *et al*. (2018) Antibody Engineering: Methods and Protocols., Methods Mol. Biol., Vol. 1827, New York, Springer New York, 269-285.

### Immobilized metal affinity chromatography (IMAC) purification

IMAC purification was carried out to remove display complexes with incompletely synthesized peptides from the reaction mixture. This was enabled by the introduction of the 6xHis tag at the C-terminus of the peptide library. The process was performed on a 200 μL scale using 160 μL of His60 Ni-magnetic beads suspension (Takara Bio, Shiga, Japan). The beads were washed twice with 500 μL of sterilized water and 500 μL of Equilibration buffer (20 mM Tris-HCl pH 7.4, 0.5 M NaCl, 20 mM Imidazole). Beads were then mixed with 40 μL of collected cDNA display supernatant and incubated at 25°C for 2 h on a rotator. The beads with bound display complexes were washed twice with 100 μL Equilibration buffer and incubated in 40 μL of Elution buffer (20 mM Tris-HCl pH 7.4, 0.5 M NaCl, 500 mM Imidazole) at 25°C for 15 min. The eluted reaction mixture was subsequently used in the selection process.

### Selection of substrate peptides for TG1

TG1 activity-based selection was performed on the 100-μL scale, using 30 μL of cDNA display solution per reaction. The reaction mixture was composed of 5 mM biotin-pentylamine (Thermo Fisher) as the acyl acceptor substrate, 10 mM Tris–HCl pH 8.0, 15 mM CaCl_2_, 5 mM DTT, and 0.95 μg/mL TG1 diluted in TG1 dilution buffer (TBS, 1% BSA, 1mM DTT). The reaction mixture was initially incubated for 60 min at 37°C during the first selection round and the time was reduced in each successive round to 20, 15, 10, and 5 min to increase the selection pressure and select the peptides with higher reactivity and specificity to TG1. In the next step, unreacted biotin-pentylamine was removed by ultrafiltration with a 3 or 10 kDa MWCO membrane (Merck Millipore). The recovered upper residual liquid (∼100 μL) was then immobilized onto the streptavidin-coated magnetic microbeads from 20 μL suspension. Immobilized beads were separated from the supernatant and washed three times with 1 mL of Selection buffer version 4 (50 mM Tris-HCl pH 7.4, 0.5 M NaCl, 1 mM EDTA, 0.7 % Tween20) and 1 mL of TE buffer followed by resuspension of the beads in 15 μL of TE buffer.

### Preparation for the selected DNA for the next selection round

The cDNA of input (cDNA displayed library before selection) and enriched (biotinylated library after selection) pools of both K5-SML and K5-SRL libraries after each selection round was used for PCR amplification with the primers 7 and 8 to prepare each DNA pool for verification of the identity of selected DNA by Sanger sequencing. After confirmation of the correct DNA backbone, another round of PCR with primers 5 and 6 was done to add the T7 promoter to DNA libraries and prepare them for the successive selection round.

### Analysis of the reactivity of enriched libraries

A TG1 enzyme reactivity assay was performed for the enriched peptide libraries from K5-SML and K5-SRL after each selection round. Due to the low molecular weight of the peptides, the peptide library was expressed as peptide-sfGFP fusion and used for the assay.

PCR amplified and purified enriched DNA pools were used as templates in a next PCR with primers 9 and 10 to obtain peptide gene fragments suitable for assembly with sfGFP gene. pRSET-K5-sfGFP plasmid (preparation explained in Supplementary material) was used as the template for PCR amplification with primers 11 and 12 to obtain the sfGFP portion for the assembly with the peptide genes. Purified PCR product after *Dpn*I digestion was used in further processing. The DNA fragments encoding enriched peptide genes were mixed with sfGFP gene portion and assembled by 1 hour incubation at 50°C using HiFi Master mix (NEB Japan, Tokyo, Japan). The reaction mixture was then used in PCR amplification with primers 13 and 14 to obtain peptide-sfGFP gene fragments suitable for cell-free protein synthesis. The reaction was done on a 5-μL scale with 25 ng of purified peptide-sfGFP DNA templates of each enriched library with incubation at 37°C for 3 h. The cell-free products (5μL) were used without further processing as substrates for TG1 reaction in a 12 μL reaction mixture composed of 2.5 mM biotin-pentylamine, 10 mM Tris–HCl pH 8.0, 15 mM CaCl_2_, 5 mM DTT, and 0.95 μg/mL TG1 for 15 min at 37°C. Quenching of the reaction was done using 2 μL of 0.5 M EDTA. Reacted samples were used in SDS-PAGE. Electrophoresis was followed by blotting of the samples onto a nitrocellulose membrane, which was then immersed in the skim milk (Nacalai Tesque, Kyoto, Japan) solution overnight at 4°C. The membrane was washed three times with PBST (Phosphate Buffered Saline, pH 7.4, 0.05% Tween 20) and incubated with Streptavidin-HRP (Invitrogen) in PBS (Phosphate Buffered Saline, pH 7.4), at 37°C for 1 hour to detect biotinylated TG1 reaction products. The membrane was washed three times with PBST and bands were detected using chemiluminescence of Amersham ECL Prime (Cytiva) HRP substrate. Detection of bands was performed using an LED light imager and imaging software MISVS II. The band densities of the image were analyzed using ImageJ software and compared in brightness to detect the highest signal corresponding to highest bulk reactivity of the enriched peptide pool.

### Analysis of the enriched DNA pools by next-generation sequencing

PCR-amplified cDNA pool of the enriched displayed library with the highest reactivity in TG1 assay was prepared for NGS analysis and analyzed using 150 bp single-read next-generation sequencing (Illumina, NextSeq550) carried out at the Center for Gene Research of Nagoya University, Japan.

### Bioinformatics analysis

First, the quality of NGS data was assessed using SeqKit (Shen et al.,2016). The probability of incorrect base identification is revealed by the quality scores for Q20 and Q30 ranges. The quality threshold was set to Q20 using SeqKit and sequences that passed the criteria were selected for further analysis. Each step was accompanied by testing the statistics of the analysis to ensure sufficient data are retained for the next step. In the following, only the DNA sequences corresponding to the peptide genes were taken and converted into amino acid sequences using Bio-python and R. To exclude unexpected outcomes, the sequences that did not contain Gln at the third position or had stop codons inside the peptide sequence were removed from the data. The enrichment factor of each peptide was calculated using the normalized count number of each peptide in input and enriched pools and was used for the creation of the heatmaps for each library to reveal preferred as well as non-preferred amino acids in each mutated position of the peptide backbone.

A hyperparamter-tuned random forest model was fitted to the data using 10-fold cross-validation and permutation importance was used to evaluate how important certain sequences were at certain positions for positive or negative peptide enrichment. During permutation importance, each amino acid position was sequentially shuffled randomly for 1000 times, and the impact on the models predictive strength was observed. In simple words, this allows to infer which amino acid patterns are critical to explain the substrate preference of TG1.

An in-house bioinformatics program was used to search for natural protein targets of TG1 in a human protein database. The database was created by extracting information from public databases and included information of about 500,000 human proteins (name, sequence, database code, etc.). Peptide matches were scored using a position weight matrix, which was calculated from the per amino acid enrichment factors of the measured TG1 substrates.

### Mini-screening of the selected peptides’ reactivity with TG1

In the mini-screening, the reactivity of top enriched peptide sequences from both libraries was compared with that of the K5 sequence to identify individual substrate peptides with a possible higher reactivity with TG1. First, the list of peptide sequences from both K5-SML and K5-SRL libraries was separately arranged according to their enrichment factor and count number. The enrichment factor, count number, and the availability of natural TG1 target motifs in peptide sequences were used as a parameter to choose the peptides for the mini-screening. The DNA of chosen peptide sequences representing peptides from both K5-SML and K5-SRL libraries were ordered from Eurofins genomics (Tokyo, Japan) as oligonucleotides and prepared for the assay as previously described. The individual peptides were expressed as fusion proteins with sfGFP in cell-free protein synthesis on a 5-μL scale and their expression was analyzed using western blotting with Anti-His-tag mAb-HRP-DirecT (MBL, Tokyo, Japan). The peptides were then checked for TG1 reactivity, with biotinylated TG1 reaction products detected on the membrane using Streptavidin-HRP. The reactivity of the individual peptides was calculated as a ratio between their respective reactivity (signal detected by Streptavidin-HRP) and expression level (signal detected by western blotting). This ratio was used for comparison of the peptide reactivity with that of the K5 sequence.

### Reactivity and cross-reactivity of the selected peptides measured by the plate assay

The reactivity of peptides selected in the mini-screening was finally verified by the TG plate assay as described before (Hitomi et al., 2009) using chemically synthesized peptides with N-terminal biotin. The plate was coated with 1 mg/mL β-casein in TBS for 30 min at 37°C. After removal of the residual casein, the wells were blocked with wash buffer (TBS, 0.1 % Tween 20) for 30 min at 37°C, followed by washing with TBS buffer. 2×peptide solution (40 mM TRIS–HCl pH 8.0, 280 mM NaCl, 30 mM CaCl2, 15 mM DTT, and the biotinylated peptides at the final concentration of 0 μM, 2 μM, 5 μM and 10 μM was added at 50 μL/well, followed by 2×TG1 solution (0.2 ng/µL of TG1 in TBS with 1 mM DTT and 1% BSA) at 50 μL/well. Cross-reactivity was checked for the peptides highly reactive with TG1, by the addition of the same concentration of TG3 enzyme and 0.15 ng/µL of TG2 enzyme diluted in TBS with 1 mM DTT and 1% BSA. The reactions were incubated for 20 min at 37°C. The quenching was done with 100 mM EDTA in TBS, and this was followed by washing with the wash buffer. The streptavidin-HRP solution diluted as 1:10000 in PBS was added at 100 μL/well. The reactions were incubated for 60 min at 37°C. The wells were washed with wash buffer and TBS. TMB (Invitrogen) was added at 100 μL/well, followed by 1 N H_2_SO_4_ at 100 μL/well after the blue color appeared. Absorbance was measured at 450 nm after the development of the yellow color (30–60 s after the addition of H_2_SO_4_). The biotinylated peptides, K5, T26 (specific substrate of TG2, Sugimura et al., 2006), and E51 (specific substrate of TG3, Yamane et al., 2010) were used as positive controls and the biotinylated peptide K5QN (Sugimura et al., 2006) with no Gln residues in the backbone was used as the negative control in the assay.

## RESULTS AND DISCUSSION

We have previously developed a cDNA display-based platform for TG substrate profiling and applied it to the analysis of TG2 substrate preference (Damnjanović et al., 2022). The same platform with modifications, namely the library design, addition of IMAC purification step for the improvement of input library quality, number of selection rounds, and refinement of bioinformatic analysis, was used in the current study to reveal the substrate preference of TG1.

Since the K5 peptide selected using the phage display technique (Sugimura et al., 2008) has a conserved portion of the sequence, shared with other peptides selected in the same study, the libraries used in the current study were designed to keep the conserved portion intact and contain random mutations in the non-conserved positions of the K5 backbone.

### DNA recovery after TG1 activity-based selection

The preparation of mRNA-linker and mRNA display libraries was confirmed by Urea PAGE (Figure 2A) and Urea SDS-PAGE (Figure 2B) respectively. mRNA displayed libraries were used for the preparation of the cDNA displayed libraries and TG1 activity-based selection, followed by the PCR amplification and gel purification of input and enriched libraries (Figure 2C). After confirmation of the correct backbone sequence by Sanger sequencing, the enriched pools of all the selection rounds were used for the evaluation of reactivity with TG1.

**Figure 2.**
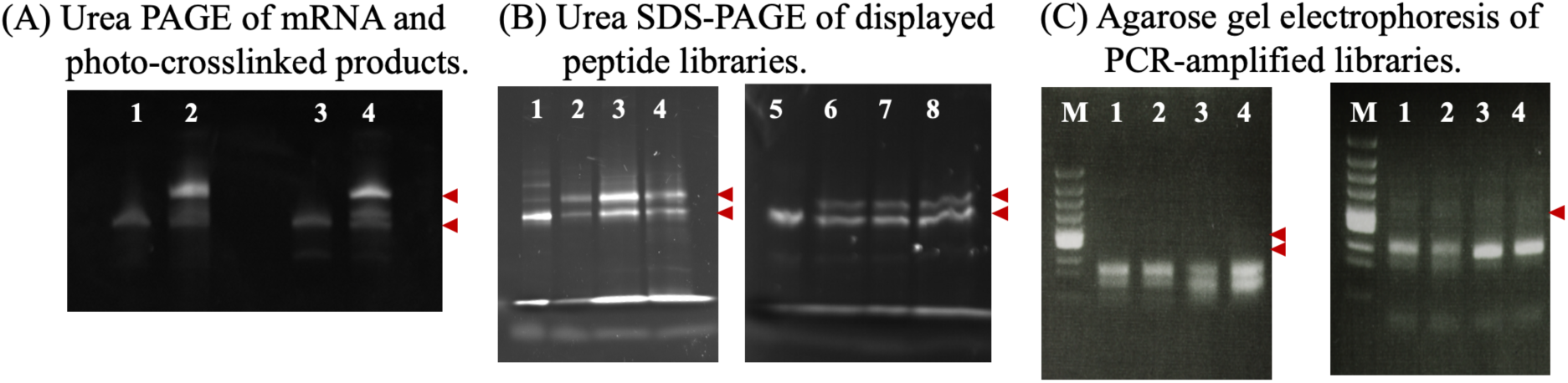
Progress of cDNA display preparation and post-selection recovery of DNA. (A): Urea PAGE of the photo-crosslinked products and free mRNA detected by SYBR Gold staining. Lanes: 1 - mRNA (K5-SML); 2 - Photo-crosslinked products of the K5-SML; 3 - mRNA (K5-SRL); 4 - Photo-crosslinked products of the K5-SRL. Above: mRNA-linker, Below: Free mRNA. (B): Urea SDS-PAGE of the *in vitro* translation products in comparison with mRNA-linker product. Lanes: 1 - mRNA-linker (K5-SML); 2 to 4 - *in vitro* translation products (K5-SML); 5 - mRNA-linker (K5-SRL); 6 to 8 - *in vitro* translation products (K5 SRL). Above: mRNA display, Below: mRNA-linker. (C): Left, Agarose gel electrophoresis of PCR amplified cDNA of the libraries. Above: PCR-amplified libraries (expected size 178 bp), Below: PCR side-product. Right, Agarose gel electrophoresis of the PCR products after gel purification. Lanes: M-φX174-*Hinc*II digest; 1 - K5-SML input library; 2 - K5-SML enriched library; 3 - K5-SRL input library; 4 - K5-SRL enriched library (expected size 178bp).

### Reactivity of enriched libraries with TG1

The reactivities of enriched K5-SML and K5-SRL libraries were evaluated using the TG1 enzymatic assay by estimating the amount of biotin incorporated into the enzymatic reaction product. For the K5-SRL library, the signal intensity of the band corresponding to the biotinylated product was found to be increasing from selection round 1 to 5 and decreased in round 6. Similarly, in the K5-SML library, the signal intensity was found to be increasing up to selection round 3 and decreased in round 4 (Figure S2). These results were used to choose the enriched libraries to be sequenced by NGS, which are the enriched pool after round 5 and its input (enriched pool after round 4) for the K5-SRL library, and the enriched pool after round 3 and its input (enriched pool after round 2) for the K5-SML library.

### Analysis of sequencing data

#### Analysis of sequencing data from K5-SML library

The heatmap (Fig. 3) revealed preferred and non-preferred residues at −1, +1, +3, +4, +5, and +6 positions from the reactive Gln. At position −1, the polar, medium-sized residue His was preferred indicating a similarity to the natural TG1 targets such as filaggrin and loricrin, followed by Gln with a similar enrichment factor. The preference for Gln-Gln motif is also observed in six out of the fourteen currently known TG1 targets such as filaggrin, involucrin, loricrin, and small proline-rich proteins 1,2 and 3. Several large hydrophobic residues were identified as the least preferred residues at −1. Pro at both, −1 and +1, positions was identified as a non-preferred feature for the enrichment in accordance with the previous study (Pastor et al., 1999). Nonpolar, and hydrophobic residues such as Trp, Ile and Phe were preferred for position +1, while Met, Gln, and several positively charged residues as Arg, His, and Lys were non-preferred residues, in agreement with the previous research (Pastor et al., 1999). Pro was preferred at the consecutive positions +4, +5 and +6. This preference could change the orientation of the peptide possibly by conferring a curved conformation which can affect the binding of the peptide with TG1. Several hydrophobic residues were not preferred at positions +4 to +6. Even though +9 is the furthest from the reactive Gln residue it was identified to have a preference for several residues, unlike position −2 which is closer to the reactive Gln but did not show any pronounced sequence preference. Position +9 is showing less preference over most hydrophobic amino acids such as Met, Cys, and Leu while preferring a range of other amino acids.

**Figure 3.**
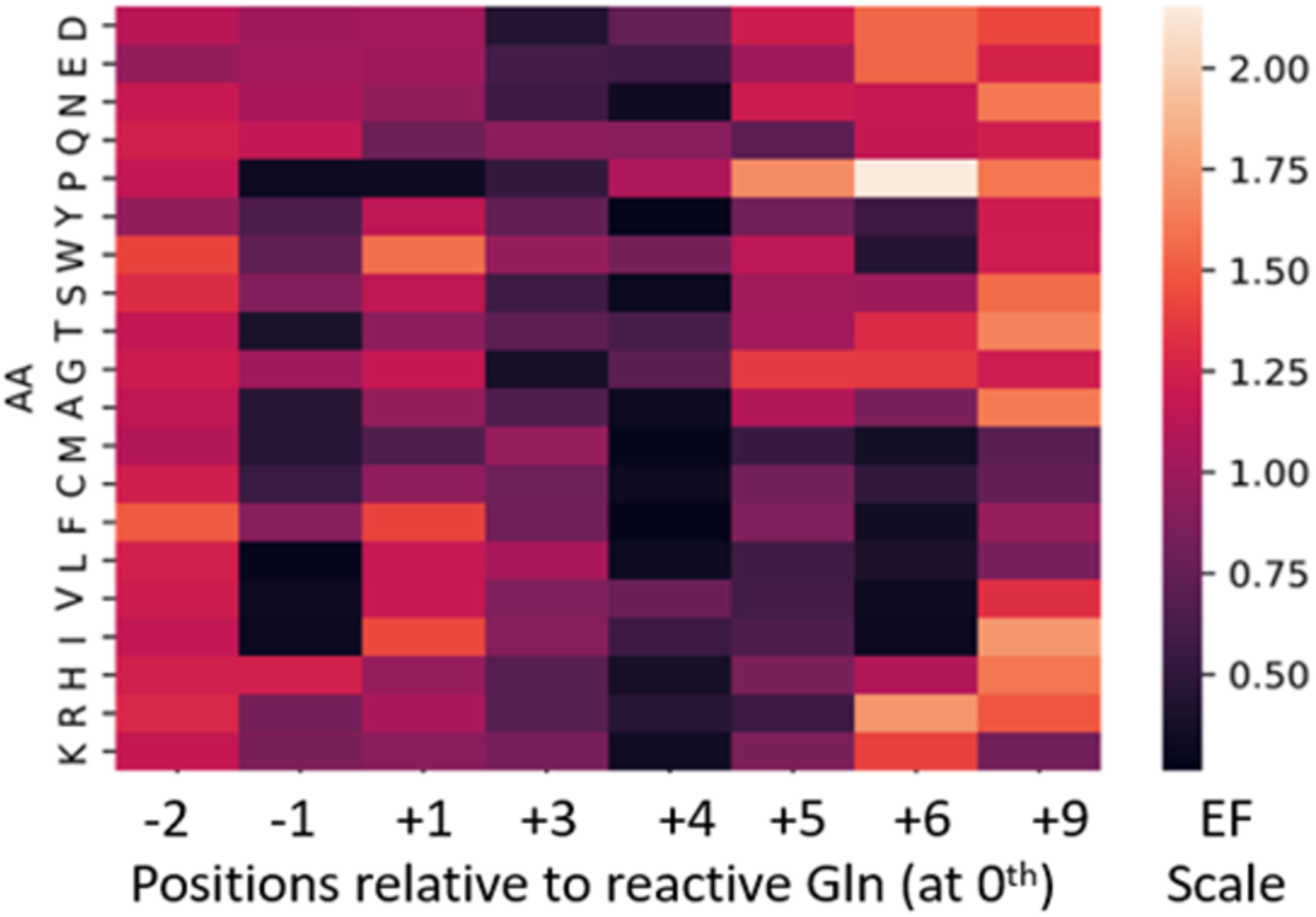
A heatmap showing per amino acid enrichment factors in the K5-SML library after the 3^rd^ selection round.

According to the permutation importance analysis (Fig. S3), the highest importance for positive enrichment was assigned to Pro at +4 followed by Leu at +3, Glu at −1, and Ser at +5. These were aligned with the amino acids contained in the K5 peptide. However, even though K5 peptide is having Phe at +9, Tyr at −2, His at +1, and Ser at +6, they were all identified as highly important for negative enrichment by the permutation importance analysis.

#### Analysis of sequencing data from K5-SRL library

The first selection round for K5-SRL library was performed and analyzed by NGS in a duplicate, with the duplicate experiments carried out under identical conditions by two different team members. The results (Fig. S4) showed that our selection method is robust and repeatable, since the pattern of per amino acid enrichment factor did not change between the two repeats. This confirmed that the selected sequences were not the result of random enrichment, but a result of properly designed and executed selection step.

To monitor the progress of the selection between different rounds and investigate whether there is a change in the residues identified as preferred or non-preferred between different selection rounds, we compared per amino acid enrichment factors of round 1, round 3, and round 5 (Fig. S5). As expected, the heatmap pattern did not change in terms of the residues identified as preferred or non-preferred from round 1 to round 5. However, by round 5, we could distinguish different level of preference that the enzyme shows to certain sequences. For example, preference at positions +5 and +6 was obvious after selection round 1 or 3, but it became pronounced after the fifth round where Ser had much higher enrichment than other amino acid sequences.

According to the permutation importance analysis (Fig. 4), the highest importance for positive enrichment has been assigned to Pro at position +4 similar to the K5-SML library. This result is well aligned with the heatmap-derived sequences and with the top 100 enriched sequences of both libraries. It is followed by Glu at position −1 and Phe at position +9, similar to the K5 peptide backbone. The highest importance for negative enrichment is assigned to Tyr at position −1 followed by Leu and Ile at position −2, Arg at +6, and His at position −1. Overall, the selected sequences from the K5-SRL library showed a consensus sequence corresponding to that of the K5 peptide, as seen by per-amino acid enrichment factors and by permutation importance analysis, indicating K5 as a specific substrate for TG1. Even though the K5 sequence can still be improved, sometimes several non-consecutive mutations in the backbone could disrupt the peptide conformation, resulting in a loss of recognition or binding affinity with the enzyme. Consequently, the selection from the two libraries used in this work favors sequences that retain the input structural features corresponding to the K5 sequence. However, when the enrichments of both libraries were considered, preferences for positions +4 for Pro, - 1 for Glu and +9 for Phe seem to be of high importance for the enrichment.

**Figure 4.**
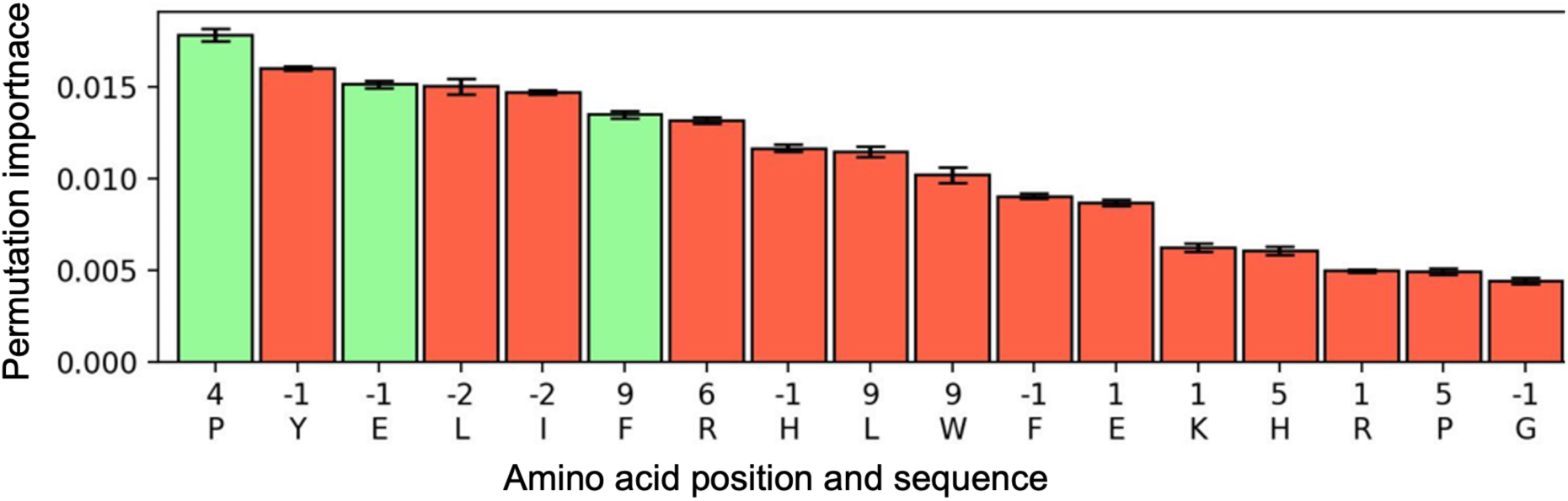
Analysis of the enriched peptides from K5-SRL. Permutation importance for the amino acids used as input for a Random Forest Regression model, sorted by importance. The permutation importance indicates how important given amino acid is to accurately predict enrichment. Color of bars indicates association with enrichment: Green = increase of enrichment, red = decrease of enrichment. Bars are mean permutation importance for ten-fold cross-validated calculations.

#### Analysis of top enriched peptide sequences selected from both libraries

We further listed the top 100 enriched peptide sequences of both libraries ranked according to the enrichment factor (Table S2). The sequence logo of the top 100 peptides selected from K5-SRL is shown in Figure 5. At position −2, Leu was preferred by 25% of the sequences followed by a 14% preference for similar hydrophobic, non-polar, and aliphatic residue, Ile, which is a different preference from that of the K5 peptide. Nearly 15% of sequences preferred Glu at position −1, similar to the K5 sequence. however, Tyr appeared in 13% of the sequences followed by Ser in 12%, and Phe and His in 11%. Preferences at the N-terminus of the peptides are aligned with the observations of the study which identified the K5 sequence (Sugimura et al., 2008), where several peptides with high reactivity to TG1 contained Tyr and Glu at the N-terminus.

**Figure 5.**
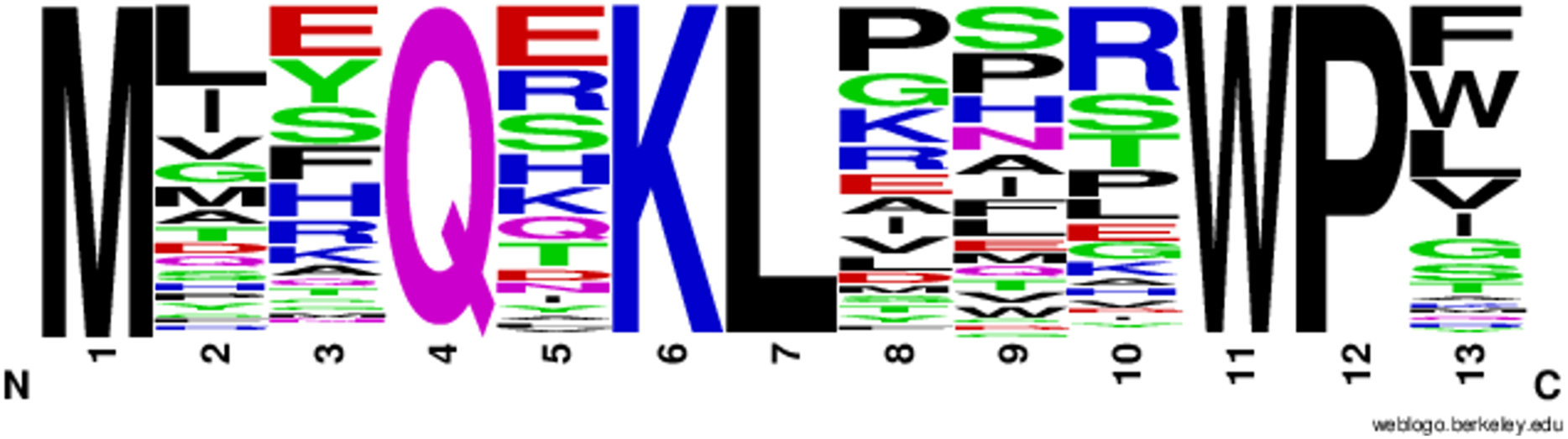
Weblogo showing the sequence of the top 100 selected peptides from K5-SRL. N-terminal Met is a formyl-methionine (from start codon) left uncleaved during the *in vitro* translation.

It was interesting to see Glu being preferred in 18% followed by Arg in 13% and Ser in 12% of peptides at position +1, even though the K5 peptide has His at this position. It seems that other charged hydrophilic residues besides His are also preferred at position +1. A portion of 14% seems to prefer Ser at the position +5 with 12% showing a preference for Pro. At position +6, Arg was preferred by more than 25% of sequences. It seems that the backbone prefers a large positively charged residue at this position over a polar one such as Ser or Thr. For position +9, both Phe and Trp are preferred by 18% of peptides, and Leu is preferred by 15%. Since the K5 peptide contains Phe at +9 position it seems that hydrophobic and non-polar residues are the most preferred for position +9. It is interesting that most of these preferences were in agreement with the previous findings on TG1 substrate preferences identified by phage display screening (Sugimura et al., 2008).

### Identification of substrate peptides having high reactivity with TG1

We analyzed the enriched peptide sequences from both libraries to identify TG1-specific substrate sequences. However, a number of peptides from K5-SML and K5-SRL had unexpected changes in the sequence such as mutations in the conserved positions of the K5 backbone. Since the quality of the NGS data was found to be good, these unexpected mutations are likely due to technical errors in DNA synthesis. As the reactive Gln was found at the expected position, we chose to include these peptides in the ranking. Eighteen peptide sequences (Table S3) were chosen and evaluated in the mini-screening as outlined in the methods. For ease of operation, the assay was conducted by separating the peptides into two sets, set 1 and set 2 (named 1-x or 2-x with x being a number to a given peptide contained in the set).

Out of eighteen peptides tested in mini-screening (Fig. S6), six most reactive ones were selected for the final evaluation in the reactions with TG1, TG2, and TG3 (Table 1) using the plate assay with the aim to identify the peptides with high reactivity to TG1 and low reactivity (cross-reactivity) to TG2 and TG3. Even though there are four TG isozymes (TG1, TG2, TG3, and TG5) identified in human skin keratinocytes or epidermal tissue (Eckert et al., 2005, 2014), we used TG2 and TG3 for the reactivity check since TG5 cannot be easily expressed in active recombinant form (Hitomi & Tatsukawa, 2015). The TRANSDAB database (http://genomics.dote.hu/mediawiki/index.php/Category: Keratinocyte_transglutaminase) was mainly used to get information regarding the TG1 substrates that have been identified so far. It was interesting to identify some of the motifs contained in the selected peptides to be present in the known TG1 substrates. Such as the sequence motif EQ in all peptides except 2-10, identified in periplakin; EQE motif present in peptide 1-2 and identified in involucrin; motifs KQP and QR present in peptide 2-7 and identified in loricrin, SPRs and desmoplakin; and presence of TQ motif in peptide 2-10 and in loricrin.

**Table 1.**
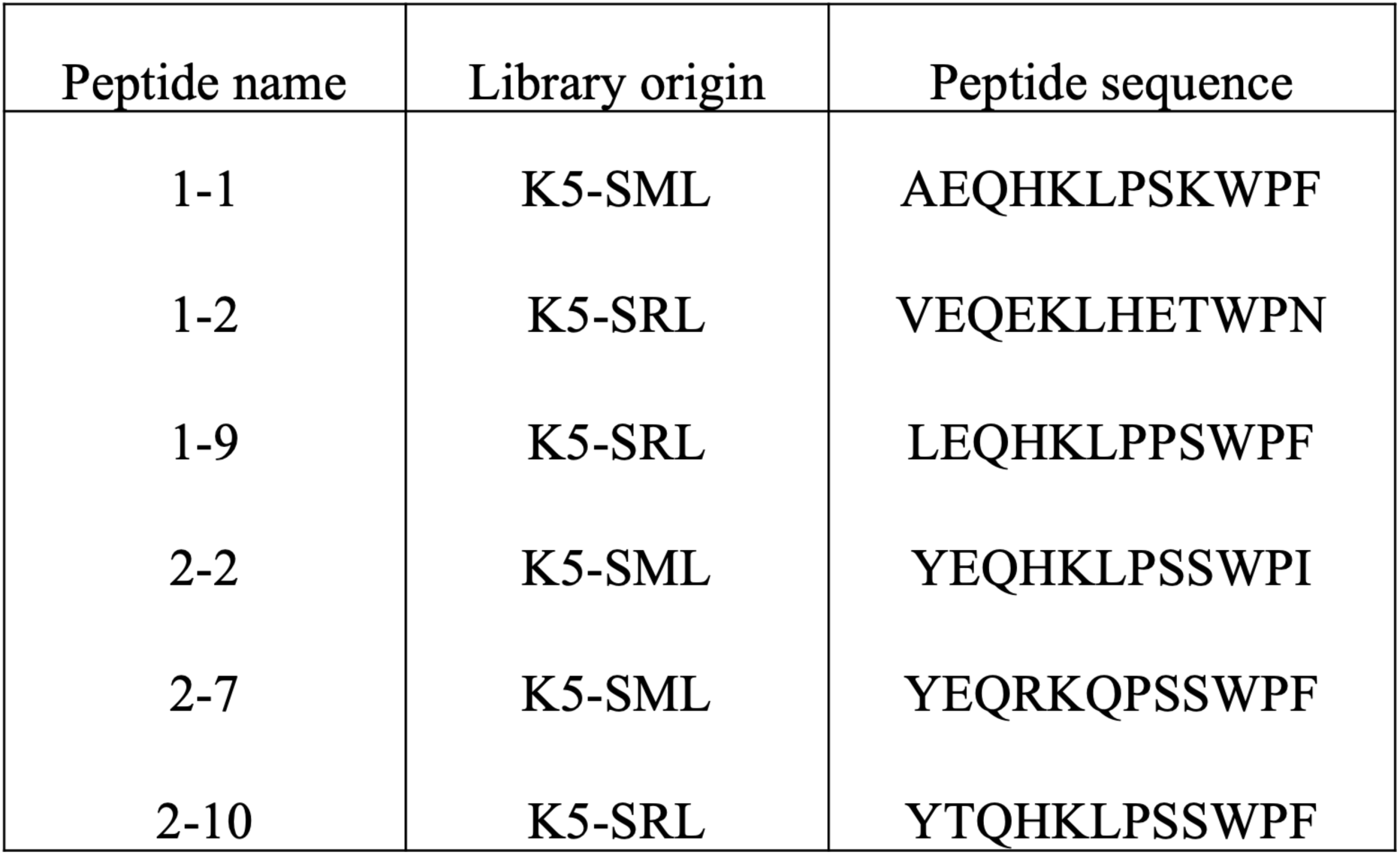
Peptide sequences chosen for verification of reactivity with TG1.

### Reactivity of the identified peptides with TG1, TG2, and TG3

The six biotinylated peptides were used for the evaluation of their reactivity with TG1 (Fig. 6). Results indicated that the reactivity of the Gln-containing peptides increased proportionally to their concentration. All selected peptide sequences were reactive with TG1. When the reactivity is compared with the K5 peptide, three peptides were identified to have a higher reactivity with TG1. Peptide 2-7 displayed the highest reactivity followed by peptides 1-1 and 2-10. The reactivity of peptides 2-7 and 1-1 was observed to be higher than that of the K5 peptide at all tested peptide concentrations. Peptide 2-10 had a similar reactivity as K5 peptide at 2 µM concentration and higher than that at 5 µM and 10 µM concentrations. Compared with the selected peptides, as expected, T26 and E51 peptides, specific substrates of TG2 and TG3 respectively, showed very low reactivity with TG1.

**Figure 6.**
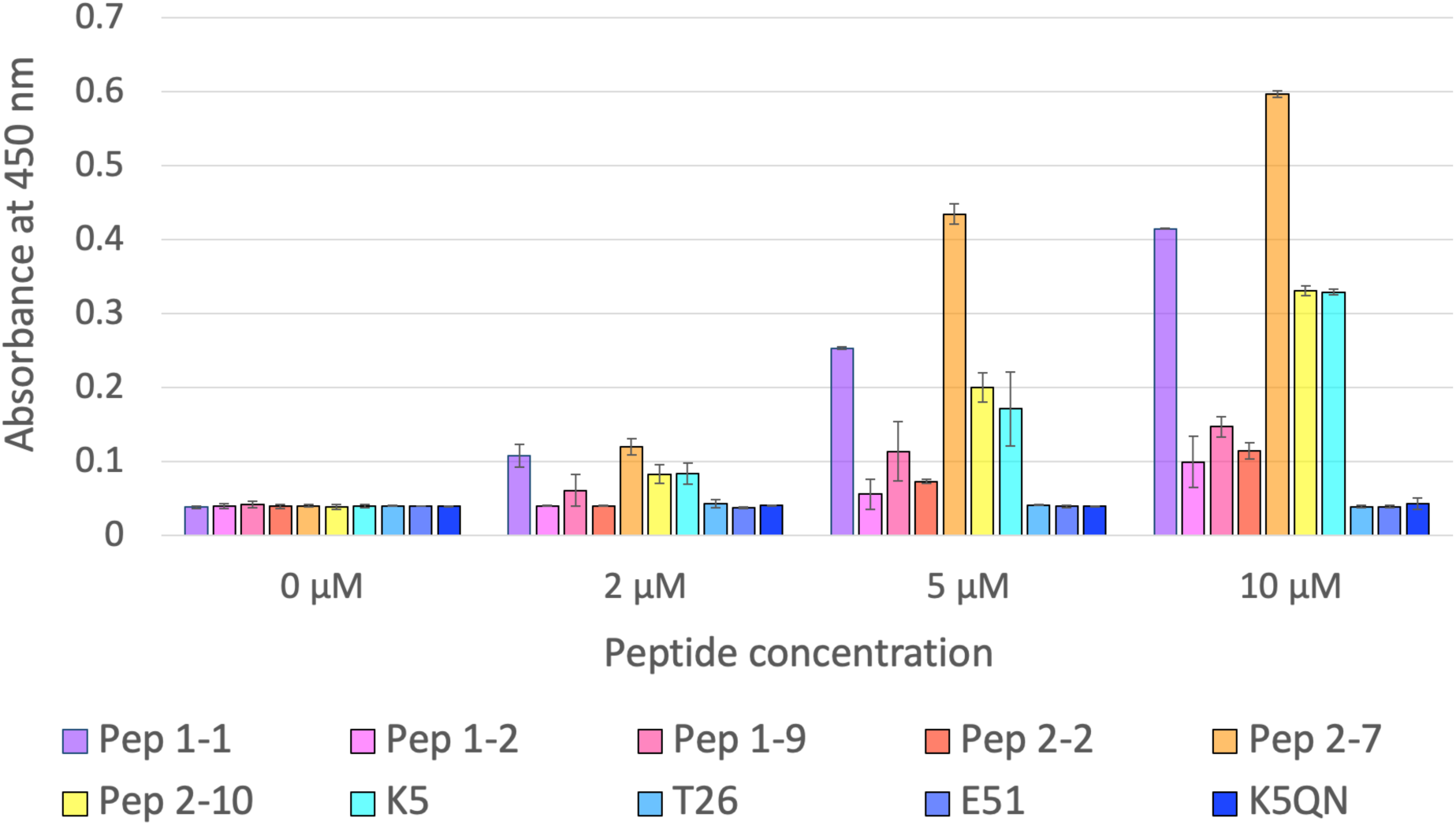
Reactivity of the peptides selected in the mini-screening in comparison with reactivities of K5 (specific substrate of TG1), T26 (specific substrate of TG2), E51 (specific substrate of TG3), and T26QN (non-substrate peptide) in the TG1 enzymatic plate assay. The peptides were chemically synthesized with N-terminal biotin for the purposes of the assay. The TG1 concentration in the assay was kept constant at all peptide concentrations. Measurement was done in duplicates with S.D. shown.

The three peptides showing the highest reactivity with TG1 were tested as TG2 and TG3 substrates (Fig. 7). In the TG2 reaction, at low peptide concentrations (2 µM), all selected peptides showed lower reactivity than the T26 peptide. The peptides 2-7 at 5 µM, and 2-10 at both 5 µM and 10µM concentrations displayed higher reactivity than the T26 peptide. Because of this, it can be considered that these two peptides are less specific as TG1 substrates. However, even at higher concentrations, peptide 1-1 was showing less reactivity with TG2 than its specific substrate T26 peptide. In terms of the cross-reactivity assay with TG3, all of the peptides showed a considerably low reactivity with TG3, when compared with the E51 peptide, a specific substrate for TG3. Peptides 1-1 and 2-10 had a similar reactivity with TG3 in all peptide concentrations. All three selected peptides, 1-1, 2-7, and 2-10, seem to be highly reactive and to some extent specific for TG1. Although peptides 2-7 and 2-10 were selected from K5-SRL, they only differed in two and one mutation respectively from the K5 sequence. This further confirmed that the K5 peptide is a very good substrate for TG1. The most specific TG1 substrate, peptide 1-1 has a sequence AEQHKLPSKWPF where −2 and +6 positions of the K5 sequence (Tyr at −2 and Ser at +6) are replaced with Ala and Lys. These mutations, especially Ser at position +6, were observed among the highly enriched peptide sequences; especially in the K5-SRL library. Since Ala is a small hydrophobic residue compared to Tyr, the mutation likely reduces the steric hindrance allowing for a better binding with the narrow active site of the TG1 enzyme. The presence of Ala at position −2 in natural TG1 targets such as desmoplakin, loricrin, and SPRs further ascertain the suitability of 1-1 peptide as a specific substrate for TG1. In terms of the other two peptides, 2-10 contains a single mutation, where Glu is replaced by Thr at position −1 adjacent to the reactive Gln. As TQ is already a known motif present in loricrin, a substrate of TG1, it is not unexpected to find it in a very reactive peptide sequence such as 2-10. Peptide 2-7 with the YEQRKQPSSWPF sequence, contains mutations at the +1 and +3 positions of the K5 backbone. At position +1, His has been replaced with Arg; a larger residue with similar chemical properties and larger size. For position +3, replacing Leu with Gln could add another reactive residue into the substrate and directly affect its reactivity with TG1. However, this hypothesis seems unlikely since the reactivity of peptide 2-7 is high with TG1, while its reactivity to TG2 or TG3 is comparable to that of the 1-1 and 2-10 peptides.

**Figure 7.**
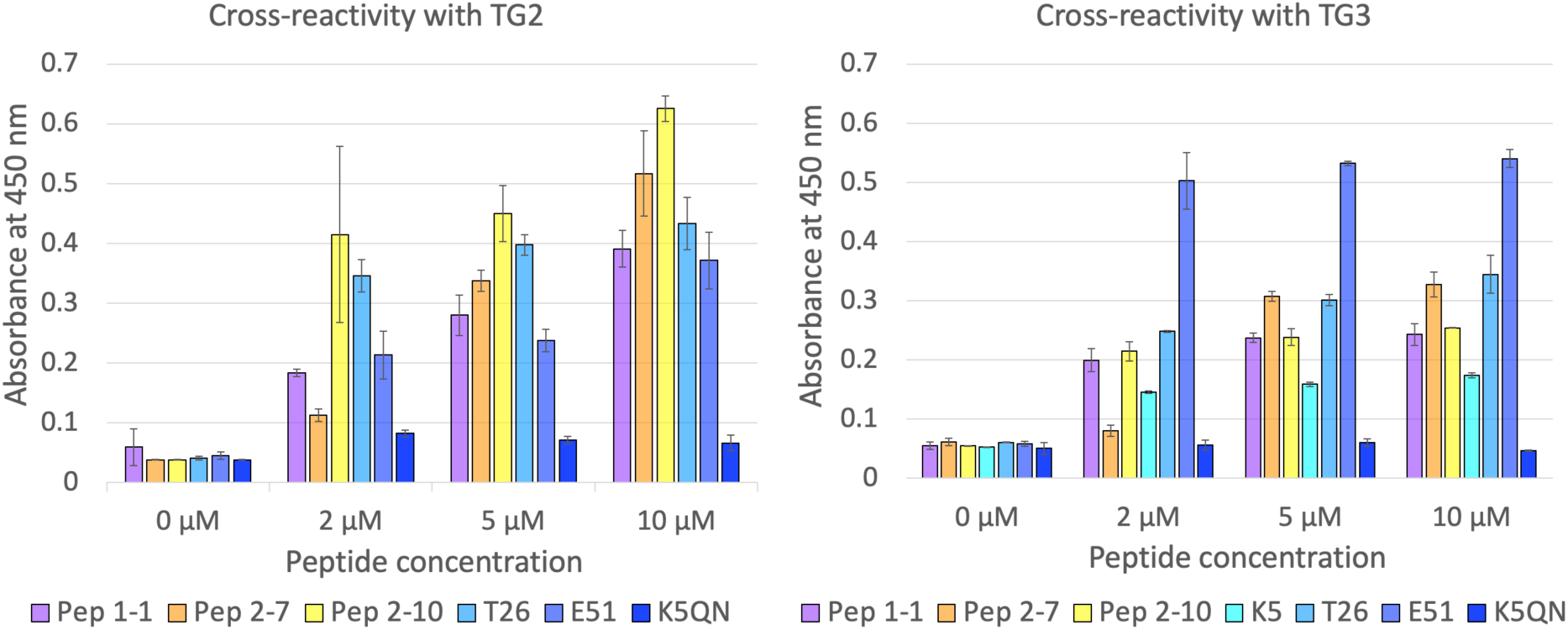
Cross-reactivity of the peptides with high reactivity to TG1 with TG2 and TG3 in comparison with reactivity of K5 (specific substrate of TG1), T26 (specific substrate of TG2), E51 (specific substrate of TG3), and T26QN (non-substrate peptide) as measured by the TG2 and TG3 enzymatic plate assays. The peptides were chemically synthesized with N-terminal biotin for the purposes of the assay. The TG2 and TG3 concentrations in the assay were kept constant at all peptide concentrations. Measurement was done in duplicates with S.D. shown.

Based on the observations, peptide 1-1 can be considered a highly reactive and specific TG1 substrate, with higher reactivity and improved specificity compared to the known K5 peptide. In fact, all three peptide sequences represent good candidates for consideration in the development of highly sensitive peptide probes for the detection of TG1, with peptide 1-1 as a leading candidate.

### Identification of potential new TG1 protein targets

The search of the human protein databases was performed using position weight matrices (PWM) of the per amino acid enrichment factors of peptides from both K5-SML and K5-SRL libraries. The results included a number of already identified TG1 targets and proteins with yet unreported relation to TG1. The Table S4 lists the top-ranking hits from both libraries which include known TG1 targets as well as other proteins.

In addition to the TG1 substrates listed in TRANSDAB, proteins such as cystatins, type II keratin, and cornifin were also identified using the PWM’s of both libraries as possible TG1 targets, which is in accordance with the previous studies (Eckert et al., 2005). When considering target proteins having a high confidence match using both PWM’s, a number of proteins were found, such as E3 ubiquitin-protein ligase, disintegrin and metalloproteinase domain containing proteins (ADAMs), zinc finger proteins, spermatogenesis-associated proteins and keratins. Even though these have not been identified as TG1 substrates, several studies have reported their possible relationship with TG1: ADAM17 is part of a cell-signaling cascade inducing TG1 expression during keratinocyte differentiation (Wolf et al., 2016). Study conducted on zinc finger protein Glis1 indicate its expression in psoriatic epidermis where TG1 expression is greatly increased, while many other zinc finger proteins play role in keratinocyte differentiation (Cassandri et al., 2017; Nakanishi et al., 2006). Similarly, studies have been conducted on the relationship between TG2 and E3 ubiquitin-protein ligase in promoting ubiquitination followed by subsequent degradation (Liu et al., 2020). A relationship among the STUB1 gene encoding E3 ubiquitin-protein ligase being regulated by TG2 has been established (Min & Chung, 2018), while its relationship with TG1 has not been reported. According to the current results, it can be certified that the bioinformatics search program can be successfully used for the identification of possible novel TG1 protein targets.

## Conclusion

In conclusion, we have identified several peptide sequences being very reactive and specific TG1 substrates, as well as amino acid patterns that have different levels of importance as preferred and non-preferred sequences. Our findings describe a detailed substrate profile of TG1 and demonstrate that the big data obtained during the peptide selection and after NGS analysis can be used to identify candidate TG1 protein targets, as candidates for further testing. We believe that the robustness and accuracy of this platform makes it a suitable tool for substrate profiling of any TG isozyme as well as any protein modifying enzyme.

## Acknowledgments

This work was supported by Grant-in-Aid for Early-Career scientists from Japan Society for the Promotion of Science (22K14828), 2022 Amano Enzyme Research Grant from Amano

Enzymes Japan, and 2023 Enzyme Research Grant from the Society for Applied Enzymology Japan, awarded to J.D. The authors are thankful to Dr. Takashi Kanamori from Gene Frontier, Japan for providing PURE*frex* components.

## Supplementary material

Supplementary material is available.

## Data availability

The data related to this article will be shared upon reasonable request to the corresponding author.

## Disclosure statement

The authors declare no conflict of interest.

## Notes

### Competing Interest Statement

The authors have declared no competing interest.

