## Supplementary material for "Substrate profiling of human Transglutaminase 1 using cDNA display and next-generation sequencing"

**Supplementary methods**

*Preparation of pRSET-K5-sfGFP plasmid*

To obtain the vector backbone, the plasmid pRSET-T26-sfGFP (Damnjanovic et al., 2022) was used as a template for PCR with primers 11 and 12 (Table S1). PCR products were confirmed by agarose gel electrophoresis, and used in further processing after *Dpn*1 digestion and column purification. pRSET-K5 plasmid was used as a template for PCR with primers 9 and 10 (Table S1) to create the insert portion. PCR products were confirmed by agarose gel electrophoresis and column-purified. In-fusion cloning was performed on a 10 μL scale using 50 ng of the insert portion (117 bp) and 100 ng of the linearized vector (3490 bp), 2 μL of 5x In-fusion cloning enzyme premix (Takara, Japan) and SW. Incubation of the mixture at 50°C for 15 minutes was followed by the transformation of DH5 alpha competent cells grown on plates with LB medium supplemented with ampicillin for 17 hours at 37°C. The identity of the clones was checked by Sanger sequencing. The colonies bearing plasmids with the correct sequence were inoculated in 4 mL of liquid LB medium containing ampicillin. Incubation for 17 hours at 37°C was followed by plasmid extraction using the standard method.

**Supplementary figures and tables**

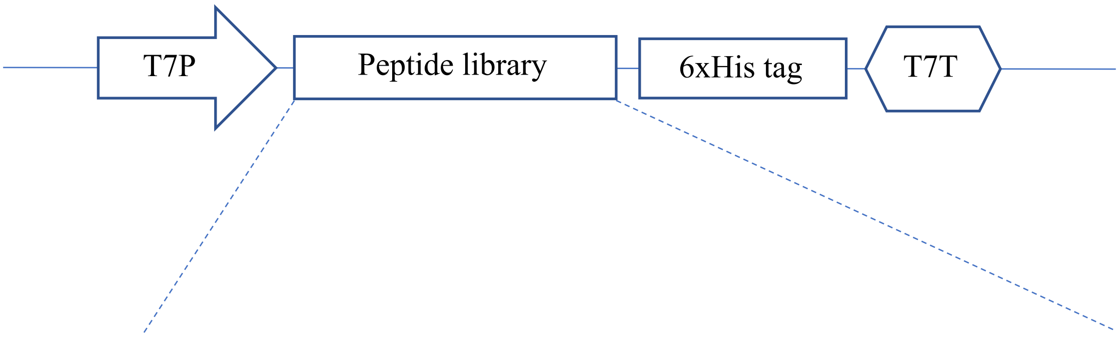


**K5:** **5'-TAT GAA CAG CAT AAA CTG CCG TCT TCT TGG CCG TTT-3'**

**K5-SML:** **5'-*NNK* GAA CAG CAT AAA CTG CCG TCT TCT TGG CCG TTT-3'**

**5'-TAT *NNK* CAG CAT AAA CTG CCG TCT TCT TGG CCG TTT-3'**

**5'-TAT GAA CAG *NNK* AAA CTG CCG TCT TCT TGG CCG TTT-3'**

**5'-TAT GAA CAG CAT AAA *NNK* CCG TCT TCT TGG CCG TTT-3'**

**5'-TAT GAA CAG CAT AAA CTG *NNK* TCT TCT TGG CCG TTT-3'**

**5'-TAT GAA CAG CAT AAA CTG CCG *NNK* TCT TGG CCG TTT-3'**

**5'-TAT GAA CAG CAT AAA CTG CCG TCT *NNK* TGG CCG TTT-3'**

**5'-TAT GAA CAG CAT AAA CTG CCG TCT TCT TGG CCG *NNK*-3'**

**K5-SRL:** **5'-*NNK NNK* CAG *NNK* AAA CTG *NNK NNK NNK* TGG CCG *NNK* -3'**

Figure S1. Schematic representation of the K5, K5-SML and K5-SRL DNA templates used for cDNA display. Gln codon is colored red. Random mutations were introduced by the NNK degenerate codon.

(A)

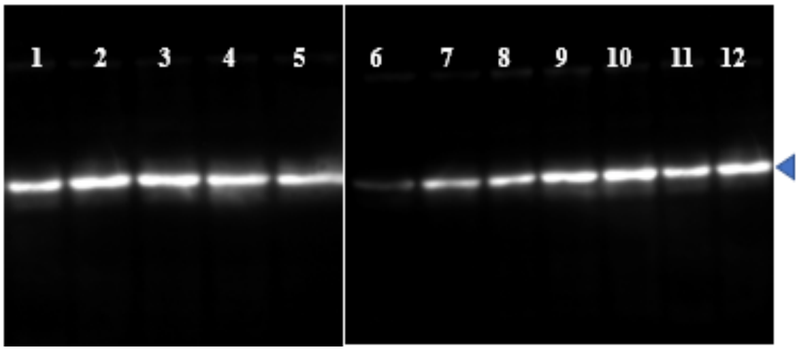


(B)

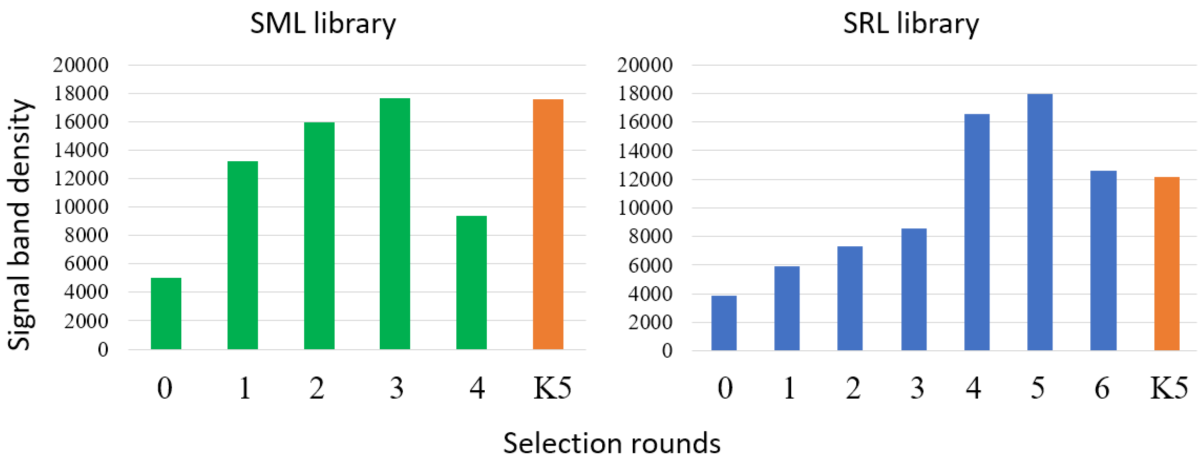


Figure S2. (A): Blotting analysis of biotinylated reaction products from the TG1-catalyzed reaction between enriched peptide libraries fused with sfGFP and pentylamine-biotin. Lanes: 1 to 4 – enriched pools from K5-SML (selection rounds 1 to 4); 5- K5-sfGFP; 6 to 11- enriched pools from K5-SRL (selection rounds 1 to 6); 12- K5-sfGFP. (B): Quantified signals from the membrane shown in (A) obtained by ImageJ. 0 refers to the original library before selection.

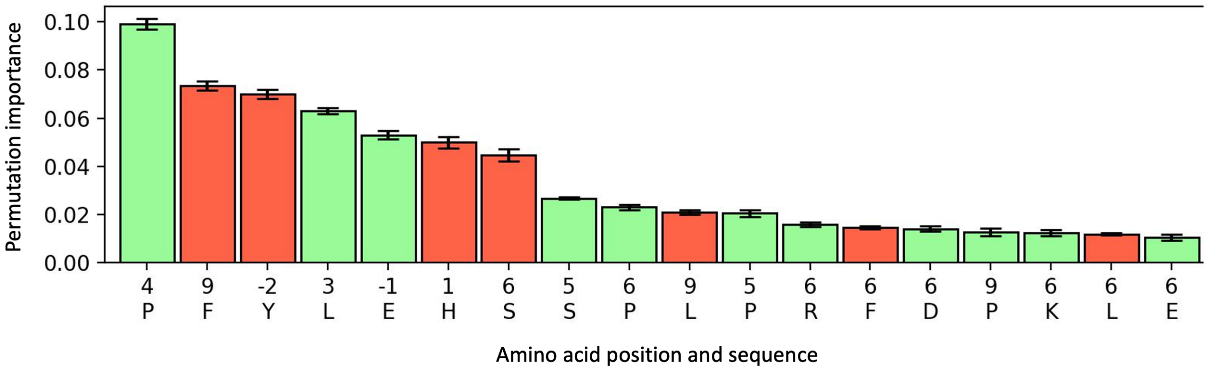


Figure S3. Analysis of the enriched peptides from K5-SML. Permutation importance for the amino acids calculated for a Random Forest Regression model, sorted by importance. The permutation importance indicates how important given amino acid is to accurately predict enrichment. Color of bars indicates association with enrichment: Green=increase of enrichment, red=decrease of enrichment. Bars are mean permutation importance for ten-fold cross-validated calculations.

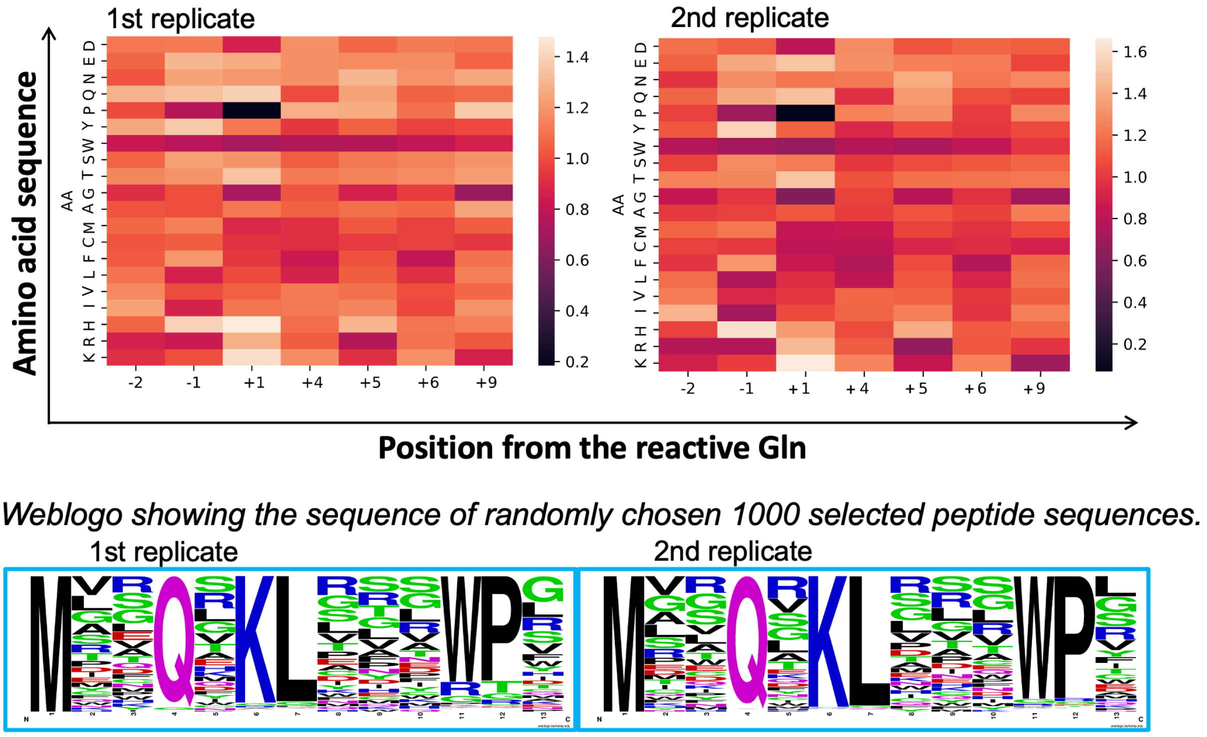


Figure S4. Analysis of the repeatability of peptide selection from K5-SRL. The first selection round was repeated under the identical conditions by two group members. The results are presented as a heatmap showing per amino acid enrichment factors at mutated positions of the peptide backbone. In the lower panel, weblogo obtained from randomly chosen 1000 selected peptides was given for both replicate selection rounds.

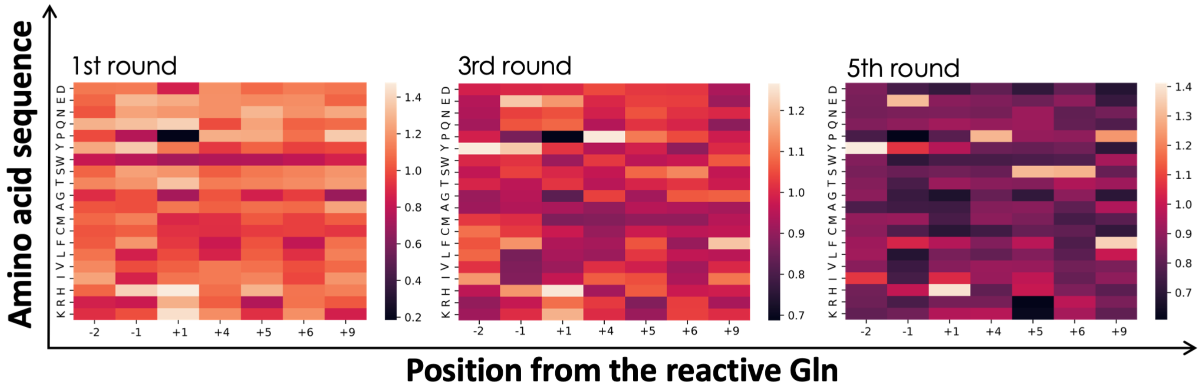


Figure S5. Analysis of the selection progress from 1$^{st}$ to 5$^{th}$ selection round for K5-SRL. The results are presented as heatmaps showing per amino acid enrichment factors at mutated positions of the peptide backbone.

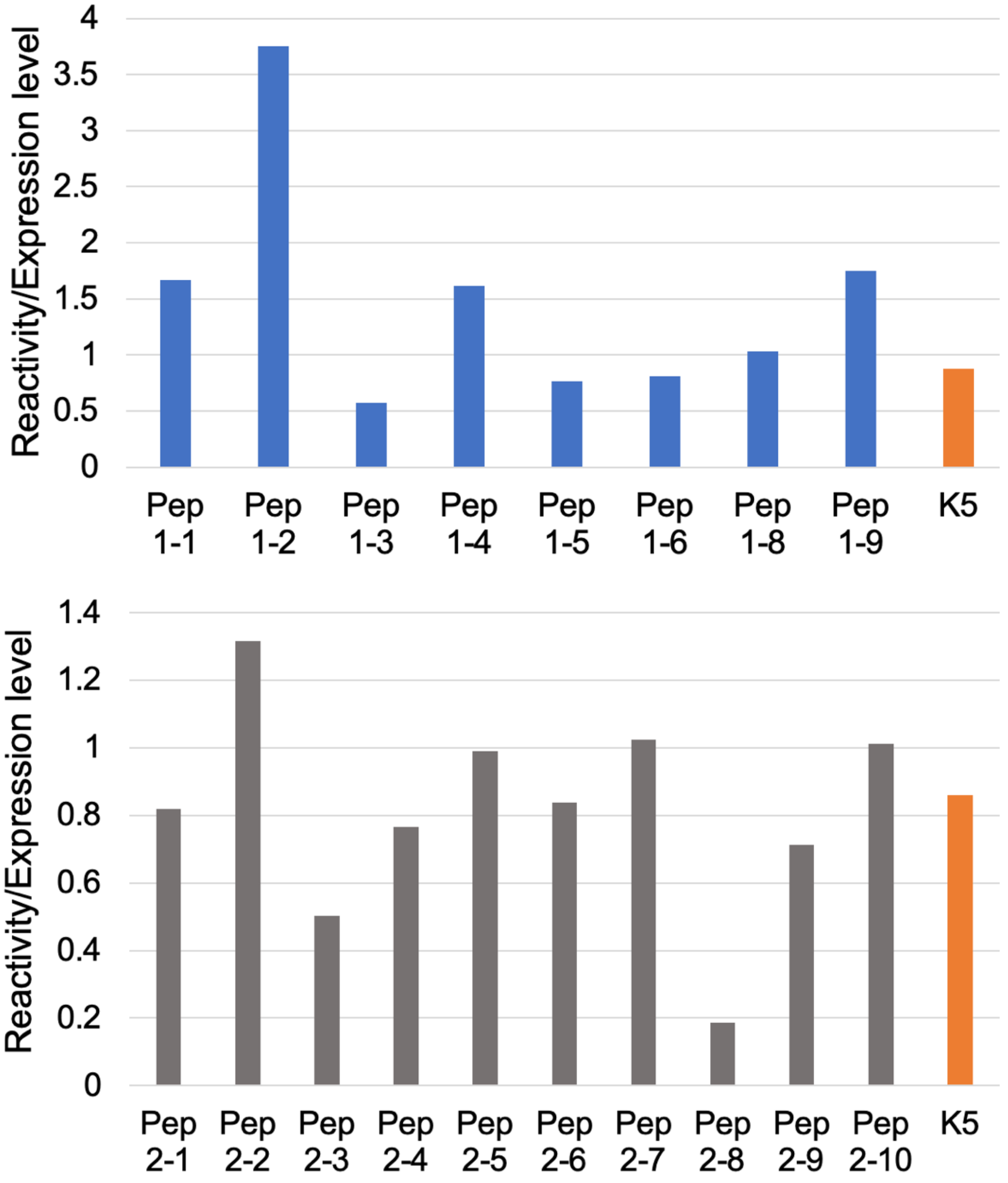


Figure S6. Peptide mini-screening using peptide-sfGFP fusion proteins obtained by cell-free protein synthesis as substrates in TG1 enzymatic assay. Y axis represents ratio of peptide reactivity and expression level detected by western blotting.

Table S1. List of primers used in this study.

| Primer number | Function | 5'-3' sequence |
|---|---|---|
| 1 | Klenow reaction | CCATGATCCAGCA |
| 2 | Preparation of pRSET for Gibson assembly | GTGGATGCTGGATCATGGTGGA |
| 3 | | TGTATATCTCCTTCTTAAAGTTAAACAAAATTAT |
| 4 | Preparation of DNA library | GATCCCGCGAAATTAATACGACTCACTATAGGG |
| 5 | | TTTCCACGCCGCCCCCCGTCCT |
| 6 | | GATCCCGCGAAATTAATACGACTCACTATAGGGAGACCACAACGGTTTCC |
| 7 | Amplification of input/enriched display | TTTCCCCGCCGCCCCC |
| 8 | | GGGAGACCACAACGGTTTCC |
| 9 | Insert amplification for assembly | GGCCGAACCTCCACC |
| 10 | | TTTCCCCGCCGCCCCC |
| 11 | Vector backbone amplification for assembly | GGTGGAGGTTCGGCCAGTAAA |
| 12 | | CCGTTGTGGTCTCCCTATAGTG |
| 13 | Obtain CFPS template | ATCTCGATCCCGCGAAATTAATACG |
| 14 | | TCCGGATATAGTTCCTCCTTTCAG |
| T7T | | GCTAGTTATTGGTCAGCGG |
| T7P | | TAATACGACTCACTATAGGG |

Table S2. The list of top 100 enriched peptide sequences from K5-SML and K5-SML.

| K5-SML Library | | K5-SRL Library | |
|---|---|---|---|
| Sequence | EF | Sequence | EF |
| MSQSKLLISWPR | 12 | VEQEKLHETWPN | 21 |
| AEQHKLPSKWPF | 11 | LSQEKLEILWPI | 20 |
| YEQNKLPSSWPP | 11 | WFQQKLSYPWPF | 20 |
| AEQQKLPSSWPF | 8 | DHQRKLINLWPW | 18 |
| EEQHKLPSDWPF | 8 | LSQRKLKDTWPF | 18 |
| VEQHKLPSSWPP | 8 | AHQQKLPHRWPL | 16 |
| YEQFTLPSSWPF | 8 | ITQEKLKSSWPT | 16 |
| YEQHKLPTSWPQ | 8 | LEQHKLPPSWPF | 16 |
| YEQRKLPSSWPA | 8 | SMQEKLVQKWPL | 16 |
| YQQHKLPSQWPF | 8 | YIQHKLPSSWPF | 15 |
| AEQHKLPSSCPF | 8 | CAQKKLKNEWPL | 15 |
| CEQHKLPSSWPT | 7 | IQQEKLKDPWPW | 15 |
| EQHKLPSGWPFG | 7 | IRQTKLKPSWPI | 15 |
| GEQHKLPSSWPK | 7 | LRQKKLGITWPW | 15 |
| HEQHKLPSSWPH | 7 | QCQWKLISSWPG | 15 |
| IEQYKLPSSWPF | 7 | VEQHKLPSSWPK | 15 |
| LEQHKLPNSWPF | 7 | VFQDKLVNRWPV | 15 |
| TEQHKLPSSWPA | 7 | VTQEKLAAPWPL | 15 |
| WEQIKLPSSWPF | 7 | VYQDKLATSWPV | 15 |
| YEQHKLPNSWPI | 7 | AYQEKLLSQWPW | 14 |
| YEQHKLPPEWPF | 7 | GFQEKLEFRWPV | 14 |
| YEQHKLPSPWPD | 7 | GHQHKLPAAWPG | 14 |
| YEQVKLPDSWPF | 7 | GYQNKLRMEWPF | 14 |
| TEQHKMPSSWPF | 7 | LEQRKLYAGWPW | 14 |
| YEQHKLPSHCPF | 7 | LKQTKLLLEWPF | 14 |
| EEQHKLPSPWPF | 6 | MNQQKLAIRWPR | 14 |
| EEQHKLPSSWPI | 6 | SEQYKLFPTWPF | 14 |
| GEQHKLPESWPF | 6 | SSQRKLPNIWPW | 14 |
| KEQHNLPSSWPF | 6 | TFQEKLRHRWPL | 14 |
| TEQHKLPSPWPF | 6 | THQFKLPSPWPT | 14 |
| TEQHKLPSSWPN | 6 | DTQRKLEVKWPW | 13 |
| WEQHRLPSSWPF | 6 | IAQIKLPGAWPT | 13 |
| YEQHELPSKWPF | 6 | IQQSKLDTYWPI | 13 |
| YEQHKKPSYWPF | 6 | LMQSKLVPVWPS | 13 |
| YEQHKLHPSWPF | 6 | LRQQKLDFTWPW | 13 |
| YEQHKLKSGWPF | 6 | MCQEKLEMRWPL | 13 |
| YEQHKLPSMWPW | 6 | SYQHKLGQRWPY | 13 |
| YEQHKLPSPWPA | 6 | VEQSKLPWRWPA | 13 |
| YEQHKMPSHWPF | 6 | VFQEKLTNLWPG | 13 |
| YEQIKLPSSSPF | 6 | YEQTKLRIEWPL | 13 |
| YEQLKTPSSWPF | 6 | YVQHKLPSSWPL | 13 |
| YEQWKLPMSWPF | 6 | AYQHKLGLRWPW | 12 |
| YNQHKLPPSWPF | 6 | CSQRKLPSSWPF | 12 |
| NEQHKHPSSWPF | 5 | HHQHKLTPHWPV | 12 |
| QEQHKLPSEWPF | 5 | HYQAKLRSYWPW | 12 |

| | | | |
|---|---|---|---|
| YEQHKLPNSWPV | 5 | IEQTKLAPGWPQ | 12 |
| YEQHKLPSPLPF | 5 | IKQTKLYETWPV | 12 |
| YEQRKQPSSWPF | 5 | LAQEKLRWRWPL | 12 |
| YHQHKLPPSWPF | 5 | LEQKKLKNTWPG | 12 |
| FEQHKLPSSWPT | 5 | LRQSKLKIGWPW | 12 |
| PEQHKLPSDWPF | 5 | LSQQKLKETWPW | 12 |
| NEQHKLPSTWPF | 5 | LSQQKLKPEWPF | 12 |
| CEQHKLPSSWPA | 5 | PFQSKLGSLWPW | 12 |
| CEQHQLPSSWPF | 5 | PKQQKLRPRWPL | 12 |
| EEQHKLPSSWFG | 5 | QGQIKLPIVWPI | 12 |
| FEQHKLPSSWPK | 5 | TSQKKLVPRWPL | 12 |
| IEQHKLPSSWPP | 5 | TYQKKLDWSWPH | 12 |
| KEQHKLESSWPF | 5 | TYQNKLPAIWPG | 12 |
| KEQHKLPSSWPA | 5 | VEQRKLGLGWPW | 12 |
| KQHKLPSSWPFG | 5 | AHQHKLPCRWPM | 11 |
| MEQHKLPHSWPF | 5 | DKQTKLMPRWPF | 11 |
| MEQHKLPPSWPF | 5 | DSQKKLKTKWPF | 11 |
| MEQHKLPSSRPF | 5 | EFQSKLGFSWPG | 11 |
| PEQHKLPSWPFG | 5 | FDQRKLFQRWPF | 11 |
| PEQHKNPSSWPF | 5 | FEQLKLVSGWPF | 11 |
| QKQHKLPSSWPF | 5 | GFQIKLPGRWPF | 11 |
| REQHKLPSSWPP | 5 | GHQKKLEVVWPV | 11 |
| YCQHKLPSHWPF | 5 | GWQEKLLHRWPF | 11 |
| YEQCKLPSSLAV | 5 | GYQLKLPSRWPV | 11 |
| YEQFKLPSSWAV | 5 | HYQEKLANLWPI | 11 |
| YEQFKLPSSWPP | 5 | IAQEKLMIRWPK | 11 |
| YEQHILPTSWPF | 5 | IHQEKLSSSWPW | 11 |
| YEQHKCPSSWAF | 5 | IKQHKLPALWPS | 11 |
| YEQHKEPSDWPF | 5 | IRQKKLSHPWPG | 11 |
| YEQHKLGSRWPF | 5 | ISQEKLDNRWPA | 11 |
| YEQHKLPASWWW | 5 | ISQNKLKHNWPV | 11 |
| YEQHKLPDSWPG | 5 | IYQDKLTMKWPS | 11 |
| YEQHKLPGSWPR | 5 | LEQRKLMWPWPS | 11 |
| YEQHKLPLSFWW | 5 | LEQRKLNVEWPL | 11 |
| YEQHKLPSEWLW | 5 | LFQEKLGFRWPI | 11 |
| YEQHKLPSFWPP | 5 | LFQRKLEQLWPI | 11 |
| YEQHKLPSIWPG | 5 | LGQSKLLHPWPL | 11 |
| YEQHKLPSKLPF | 5 | LHQSKLVLDWPY | 11 |
| YEQHKLPSLWPE | 5 | LKQSKLLPRWPL | 11 |
| YEQHKLPSNRPF | 5 | LNQSKLGLGWPV | 11 |
| YEQHKLPSSRQF | 5 | LRQAKLIMHWPW | 11 |
| YEQHKLPSSRTF | 5 | LRQEKLEAHWPI | 11 |
| YEQHKLPYNWPF | 5 | LSQYKLGSKWPF | 11 |
| YEQHKLPYSWPS | 5 | LYQDKLAHTWPW | 11 |
| YEQHKLVSSWPN | 5 | MEQTKLGFDWPF | 11 |
| YEQHKPPSEWPF | 5 | MFQNKLRETWPW | 11 |
| YEQHKVPSSLPV | 5 | MQQQKLGFRWPT | 11 |
| YEQHKWPSHWPF | 5 | MRQVKLWTRWPF | 11 |

| | | | |
|---|---|---|---|
| YEQHKWPSSLTF | 5 | MSQSKLPAFWPV | 11 |
| YEQHTLPPSWPF | 5 | NYQYKLPPAWPM | 11 |
| YEQHTLPSRWPF | 5 | PHQSKLIHRWPL | 11 |
| YEQKKLQSSWPF | 5 | QEQRKLRPRWPS | 11 |
| YEQLKLPFLAVW | 5 | QHQRKLPVTWPR | 11 |
| YEQNKLPSSRPF | 5 | RPQVGDIHMVEQ | 11 |
| YEQNKLTSGWPF | 5 | RYQKKLICPWPF | 11 |

Table S3. The peptide sequences selected for the mini-screening.

| | | DNA library | Peptide name | Amino acid sequence |
|---|---|---|---|---|
| 1 | Set 1 | K5-SML | 1-1 | AEQHKLPSKWPF |
| 2 | | K5-SRL | 1-2 | VEQEKLHETWPN |
| 3 | | | 1-3 | WFQQKLSYPWPF |
| 4 | | | 1-4 | LSQEKLEILWPI |
| 5 | | K5-SML | 1-5 | YEQHKLPSPWPF |
| 6 | | | 1-6 | YQQHKLPSQWPF |
| 7 | | | 1-8 | FEQHKQPSSWPF |
| 8 | | K5-SRL | 1-9 | LEQHKLPPSWPF |
| 9 | Set 2 | K5-SML | 2-1 | YEQWKLPSSWPV |
| 10 | | | 2-2 | YEQHKLPSSWPI |
| 11 | | | 2-3 | YEQHKLPSSWPT |
| 12 | | | 2-4 | YEQHKLQSRWPF |
| 13 | | | 2-5 | YEQHKLQSSWPV |
| 14 | | | 2-6 | YEQHKLPSRWPF |
| 15 | | | 2-7 | YEQRKQPSSWPF |
| 16 | | K5-SRL | 2-8 | YLQHKLPSSWPF |
| 17 | | | 2-9 | YEQHKLPSSWQE |
| 18 | | | 2-10 | YTQHKLPSSWPF |

Table S4. Summarized results of the search for TG1 protein targets using the in-house bioinformatics tools. The maximum score was 15.36 for the K5-SML and 14.11 for the K5-SRL. Only the top-ranking outcomes with scores exceeding 13 for K5-SML and 12 for K5-SRL were included in the analysis. Selected results are provided in the table below for reference.

| **Match** | **K5-SML library** | **Identified motifs** | **K5-SRL library** | **Identified motifs** |
|---|---|---|---|---|
| Already known TG1 targets | Desmoplakin | LNQWKT | Huntingtin interacting protein | DTQLKL |
| | Involucrin | EQQLKQ | Microtubule-associated protein | ERQRKL |
| | Huntingtin interacting protein | KDQRKM<br>NHQLKE<br>RFQRKQ<br>AKQGKM<br>VEQVKR<br>QGQRKT | Small proline-rich protein 1 | PCQPKL |
| | Microtubule-associated protein | RVQSKI<br>RKQKKR | | |
| | Small proline-rich protein 1 | PAQQKT | | |
| | Small proline-rich protein 2 | PCQDKC<br>QCQQKC | | |
| New potential targets | Connector enhancer of kinase suppressor of ras 2 | FQQWKQ | Prematurely terminated mRNA decay factor-like | IEQHKL |
| | Nucleolar transcription factor 1 | SQQWKL | E3 ubiquitin-protein ligase TTC3 | YEQIKL |
| | Ellis-van Creveld syndrome protein | RQQWKL | Tetratricopeptide-like helical domain-containing protein | YEQIKL |

| | | | | |
|---|---|---|---|---|
| | kinase anchor protein 4 | FNQWKQ | Protein bicaudal C homolog 1 | YEQKKL |
| | Calcium-binding tyrosine phosphorylation-regulated protein | FHQIKV | Radixin | LEQHKL |
| | Glycine N-acyltransferase-like protein 2 | WHQWKC | Moesin | LEQHKL |
| | Carbohydrate sulfotransferase 2 | FQQIKQ | Dynein heavy chain 6 | FEQHKL |
